# *Greb1* is required for axial elongation and segmentation in vertebrate embryos

**DOI:** 10.1101/692095

**Authors:** Ravindra Singh Prajapati, Richard Mitter, Annalisa Vezzaro, David Ish-Horowicz

**Affiliations:** Cancer Research UK Developmental Genetics Laboratory; Centre for Craniofacial Development and Regenerative Biology, King’s College London, London, SE1 9RT, UK; Francis Crick Institute, 1 Midland Rd, London NW1 1AT, UK; Veyrier, 1255, Switzerland.; Cancer Research UK Developmental Genetics Laboratory, and University College London; Medical Research Council Laboratory of Molecular Biology, University College London, Gower St, London WC1E 6BT

**Keywords:** tailbud, neural tube, axial stem cells, somites, transcriptome, progenitors, clock

## Abstract

During vertebrate embryonic development, the formation of axial structures is driven by a population of stem-like cells that reside in a region of the tailbud called the chordoneural hinge (CNH). We have compared the CNH transcriptome with those of surrounding tissues and shown that the CNH and tailbud mesoderm are transcriptionally similar, and distinct from the presomitic mesoderm. Amongst CNH-enriched genes are several that are required for axial elongation, including *Wnt3a, Cdx2, Brachyury/T* and *Fgf8*, and androgen/estrogen receptor nuclear signalling components such as *Greb1*. We show that the pattern and duration of tailbud *Greb1* expression is conserved in mouse, zebrafish, and chicken embryos, and that *Greb1* is required for axial elongation and somitogenesis in zebrafish embryos. The axial truncation phenotype of *Greb1* morphant embryos is explained by much reduced expression of *No tail* (*Ntl/Brachyury*) which is required for axial progenitor maintenance. Posterior segmentation defects in the morphants (including misexpression of genes such as *mespb, myoD* and *papC*) appear to result, in part, from lost expression of the segmentation clock gene, *her7*.

## INTRODUCTION

Vertebrate embryos develop in a highly organized fashion, progressively laying down axial tissues as they elongate along the anteroposterior embryonic axis (Brown and Storey, 2000; Catala et al., 1996; Wilson and Beddington, 1996; Wilson et al., 2009). Serial transplantation and other lineage tracing studies in mouse and chick have shown that a self-maintaining region in the tailbud called the chordoneural hinge (CNH) includes multipotent stem-cell-like progenitors for axial structures (Brown and Storey, 2000; Catala et al., 1996; Wilson and Beddington, 1996; Wilson et al., 2009). These include bipotent neuromesodermal progenitors (NMPs) that can generate both neural and mesodermal cells (Cambray and Wilson, 2002; Cambray and Wilson, 2007; McGrew et al., 2008; Selleck and Stern, 1991; Tam and Tan, 1992; Tzouanacou et al., 2009).

Adjacent to the CNH is the tailbud mesoderm (TBM) which contains the precursors of the axial mesoderm in an unsegmented tissue, the presomitic mesoderm (PSM; Fig. 1A). During elongation, the PSM is displaced posteriorly while its anterior buds off a series of somites, epithelial balls that develop into segmental mesodermal structures such as the axial skeleton and musculature (reviewed in Pourquie, 2011)

**Fig. 1.**
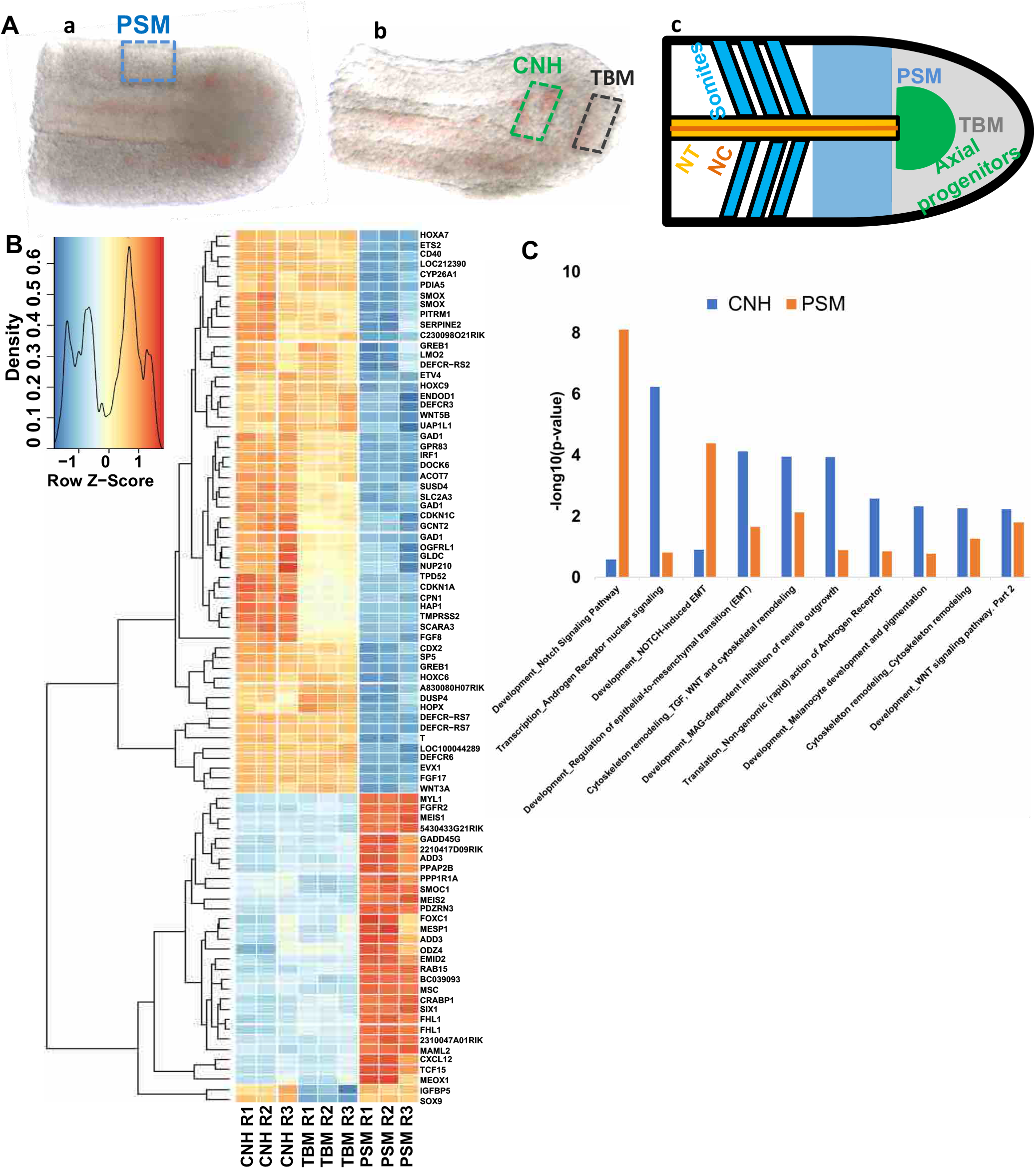
CNH transcriptome is distinct from PSM. (A) Dissection of PSM, CNH, and TBM of mouse at E10.5; (a) dorsal view of E10.5 tail, black dotted rectangle represents dissected PSM and, (b) lateral view of (a) after removing PSM from last somite till end of tail; (c) schematic of tail regions with anterior to the left. Text colours correspond to those of different posterior axial regions colours (NT: neural tube; NC: notochord; PSM: presomitic mesoderm; TBM: tail bud mesoderm). (B) Heatmap showing differentially expressed genes (Fold change >2 and <-2, and p-value <0.05) in the CNH, TBM, and PSM. (C) Pathway enrichment analysis (see Supplementary Information). y-axis shows -log_10_(p-value) with enriched GO terms along the x-axis.

Several studies have illuminated how axial progenitors are maintained during anteroposterior elongation. Briefly, a positive feedback loop between Brachyury/T and Wnt3a maintains axial progenitors in the tail bud (Martin and Kimelman, 2010; Wilson et al., 2009). In parallel, Fgf signalling protects axial progenitors from differentiation induced by Retinoic acid (RA) that is secreted by differentiating and young somites and diffuses into the PSM (Diez del Corral et al., 2003; Olivera-Martinez et al., 2012; Ribes et al., 2009).

However, Fgf8, Wnt3, and T are all expressed in much larger domains than the CNH and so do not specifically distinguish axial progenitors from more specialised cells such as the TBM. Transcriptome analysis of dissected axial progenitor tissue during the period of axial elongation and of in vitro-derived NMPs has identified genes that are differentially expressed between progenitors and presomitic mesoderm cells (Gouti et al., 2017; Olivera-Martinez et al., 2014; Wymeersch et al., 2019). However, the functional significance of many of these genes has yet to be defined.

In this paper, we explore the transcriptional profiles of the CNH, TBM and PSM of E10.5 mouse embryos. We find that the CNH transcriptome is very similar to that of the TBM, and significantly different from that of the PSM. Several genes are expressed in both the CNH and TBM but not in the PSM, although none exclusively mark the CNH. Amongst the CNH-enriched transcripts is *Greb1* which encodes a transcriptional co-activator for androgen/estrogen hormone signalling. We show that *Greb1* is expressed in the tailbud in mouse, chick and zebrafish embryos, and is required for axial progenitor maintenance and somite compartmentalisation in zebrafish. Our results indicate that Greb1 plays an evolutionarily-conserved role during vertebrate axial extension and segmentation.

## RESULTS AND DISCUSSION

### CNH transcriptome is distinct from PSM but TBM

To identify potential markers for the CNH, we used microarray analysis on dissected tissue regions to identify genes whose expression in the E10.5 mouse CNH is elevated relative to that in the PSM and TBM (Fig. 1A). 150 genes were upregulated and 98 downregulated comparing the CNH to the PSM (Table S1). Only 12 up-and 2 down-regulated transcripts distinguished the CNH and TBM, consistent with the latter population being directly derived from the former (Table S1).

To confirm that many genes identified by microarray analysis are selectively expressed in progenitor regions of the extending embryo, we searched the Mouse Genome Informatics (MGI) database (Finger et al., 2017) for the expression patterns of 53 genes whose expression were upregulated ≥2-fold in the CNH (Fig. 1B). A majority of these genes (29/53) are annotated as being expressed in tissues related to axial elongation, i.e., in one or more of the primitive streak, node, tailbud, and future spinal cord (Table S2). By contrast, most downregulated genes (23/27 reduced ≥2-fold) are expressed in more specialised progeny cells, i.e., somites, unsegmented mesoderm or neural tube (Table S2).

### Greb1 expression coincides with axial elongation in vertebrate embryos

We also compared our list of CNH-enriched genes with those previously identified in previous studies of the CNH or NMPs (Table S3; Gouti et al., 2017; Olivera-Martinez et al., 2014; Wymeersch et al., 2019). Expression of seven of the ten most-enriched genes *(Fgf8, Cdx2, T, Wnt3a, Sp5, Evx1,* and *Fgf17)* was previously reported in the CNH and TBM, and to be functionally important for axial development (Cambray and Wilson, 2007; Dunty et al., 2014; Maruoka et al., 1998; Takada et al., 1994).

The expression and roles during axial elongation and segmentation of the remaining three most CNH-enriched genes from our study *[Defcr-rs7, Defer-rs6* (which encode small immune-defect peptides) and *Greb1*] have not been previously studied. We focused on *Greb1*, which encodes a co-activator of the Estrogen and Androgen receptors that is active in human estrogen-receptor-positive primary breast and prostate cancer cells (Lee et al., 2019; Mohammed et al., 2013). Androgen receptor nuclear signalling is the most CNH-enriched pathway revealed by pathway enrichment analysis of our differentially expressed genes (Fig. 1C; Table S4; Supplementary Information), and enriched *Greb1* expression has been found in previous studies of axial and neuromesodermal progenitors (Table S3; Gouti et al., 2017; Olivera-Martinez et al., 2014; Wymeersch et al., 2019).

First, we visualised *Greb1* transcription in elongating mouse embryos using *in situ* hybridisation (E10.5-E13.5; Materials and Methods). *Greb1* expression is restricted to the CNH and dorsal TBM by E10.5, (Fig. 2A,A’). Such expression is maintained during axial elongation, weakens by E12.5, and is lost at E13.5 when axial elongation ceases (Fig. 2B,B’,C,C’).

**Fig. 2.**
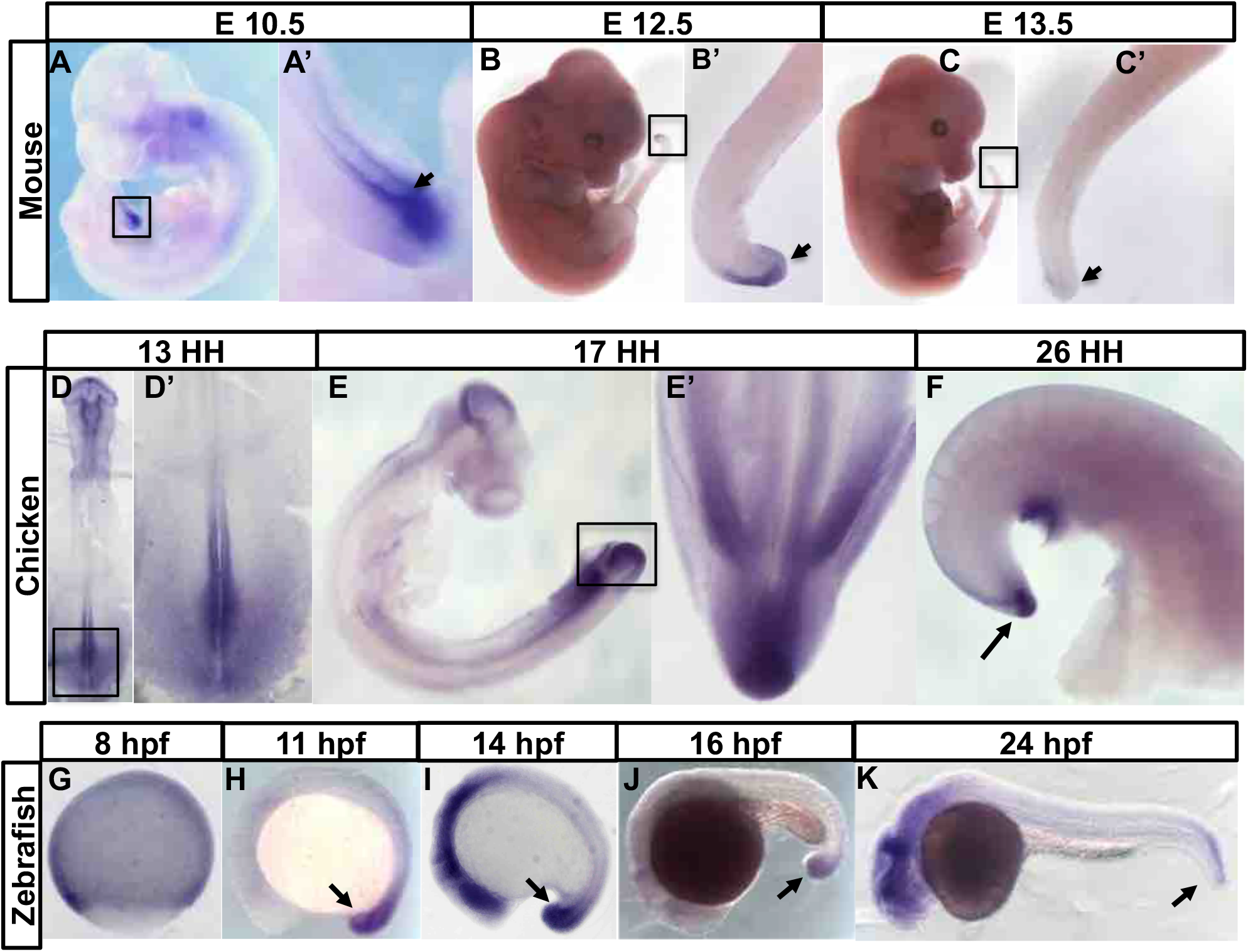
The timing of axial *Greb1* expression is coincident with axial elongation in vertebrate embryos: (A–C) Mouse embryos and their tail regions at different embryonic stages; (A,A’) Lateral view of E10.5 embryo, showing the *Greb1*-expressing tail region (boxed). (B) Lateral view of E12.5 and (B’) its tail region, showing reduced *Greb1* expression. (C) Lateral view of E13.5 and (C’) its tail region, showing that expression is lost. (D–H) *Greb1* expression in chick embryos at different stages: (D) dorsal views of HH13 stage and (D’) its tail region; (E) is lateral view at HH17 stage and (E’) its ventral view of tail region; F is a lateral view of HH26 stage. *Greb1* expression in the tailbud (arrowed) is almost gone. G-K are lateral view of zebrafish embryos of the indicated ages (tailbud region arrowed).

The above results show that, although not restricted to the CNH, axial *Greb1* transcription in early mouse embryos coincides in time and place with the processes of axial extension and segmentation. To test if this correlation is evolutionarily conserved, we examined *Greb1* expression in chick and zebrafish embryos. In both animals, *Greb1* expression in the tailbud starts during elongation, and terminates when elongation and segmentation is complete. *Greb1* is expressed in the HH13 chick caudal neural plate, whose cells contribute to the neural tube, somites, and notochord, node and primitive streak (Fig. 2D,D’). Its axial transcription then becomes confined to the region of the tailbud which includes the chick CNH and TBM (HH17; Fig. 2E,E’; McGrew et al., 2008), and has almost completely decayed when elongation is complete (HH26; Fig. 2F).

In zebrafish embryos, *Greb1* transcription becomes confined to the region of the tailbud that contains axial progenitors (Fig. 2G-K). It persists during segmentation (11-16 hpf; Fig. 2H-J), and disappears when axial elongation comes to an end (24 hpf; Fig. 2K). This conserved spatial and temporal timecourse in early vertebrate embryos strengthens the link between *Greb1* expression and axial extension.

### Knock-down of *GREB1* disrupts axial elongation

To test if *Greb1* is functionally required during elongation and segmentation, we knocked down its expression by injecting antisense morpholinos into 1-2 cell zebrafish embryos (Materials and Methods). We used two *Greb1* splicing-blocking morpholinos (M1 and M2) that target the exon2-intron2 and exon16-intron16 boundaries, respectively (Fig. S1). These oligos should interfere with mRNA splicing to cause skipping of the adjacent exon and a shifted translational reading frame. The ensuing premature translational termination would completely truncate Greb1 protein (M1) or encode one that is only 40% full-length (M2). As a control, we also injected a mismatched morpholino (MM) based on M2 but with 5 bases mutated to prevent binding to the primary *Greb1* transcript.

We verified the splice-blocking activities of both morpholinos via RT-PCR on RNA from injected embryos. *Greb1* splice variants corresponding to misprocessed transcripts were detected in M1- and M2-injected morphants but not MM morphant embryos (Fig. S1). DNA sequencing of these variant products confirmed that they result from skipping of the appropriate exons: exon 2 for oligo M1, and exon16 for M2 (Fig. S1).

We assayed the effects of *Greb1* knockdown 24 h after injection into embryos, when extension and segmentation is complete. M1 and M2 morphant embryos suffer three major axial defects: a curved trunk; a reduction in total body length (head-to-tail); misshaped somites and indistinct somite boundaries predominantly in more posterior axial regions (Fig. 3A-C). Injection of 4 ng/μl blocking oligonucleotide generates a high frequency of embryos showing all three defects (50/107 injected embryos for M1; 61/110 embryos for M2). No such abnormalities are seen in embryos injected with the control MM morpholino (0/15). Injecting 2 ng/μl of morpholino causes similar defects, albeit at lower frequencies (M1: 11/46; M2: 15/36; MM: 0/8).

**Fig. 3.**
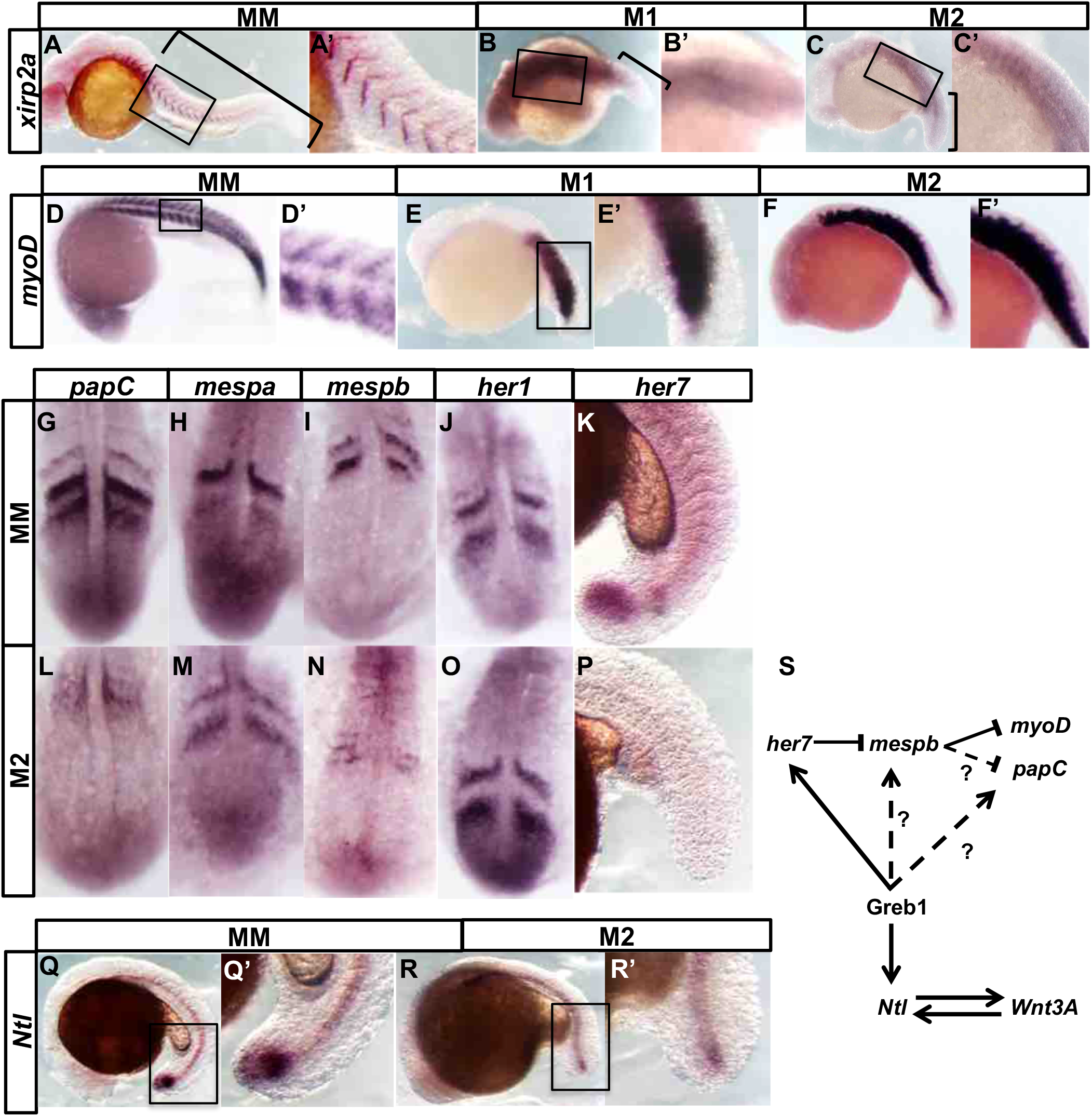
*Greb1* is required for axial elongation: (A-C, A’-C’): lateral views of zebrafish embryos at 24 hpf, showing: (A) wildtype chevrons of *xirp2a* expression and the tail region (bracketed); (B, C) posterior loss in M1 and M2 morphants. The tail regions that are truncated and contain disrupted somites are bracketed. (D-F, D’-F’): expression of *myoD* in control (MM), M1 and M2 morphants. (G, L) *papc;* (H, M) *mespa;* (I, N) *mespb;* (J&O) *her1;* (K, P) *her7;* (Q, R) and (Q’, R’) *Ntl* expression in the tail region of 15 hpf control and morphant embryos. (S) shows a tentative model for gene interactions between Greb1 and patterning genes. Anterior expression of Her7 restricts *mespb* expression to the posterior somite compartment which, in turn, restricts *myoD* and *papC* expression to the anterior compartment. Continuous arrows show interactions that are likely to be direct. Dashed arrows could be direct or indirect.

These phenotypes are not due to unspecific toxicity from the injection. We coinjected each morpholino with one that knocks down *p53* expression, thereby preventing previously reported oligo-induced p53-dependent cell death (Supplementary Methods; Robu et al., 2007). Each blocking morpholino still efficiently caused axial extension and segmentation phenotypes (M1: 20/30; M2: 25/37; control MM: 0/10). Together, our data suggest that normal axial elongation and segmentation is dependent on Greb1 activity.

### Greb1 is needed to maintain *Ntl* expression in the tailbud

The axial truncations of *Greb1* morphants resembles the phenotype of embryos mutant for *No tail (Ntl),* the zebrafish homologue of *Brachyury/T,* which is expressed in the tailbud, posterior PSM and notochord of wildtype embryos (Halpern et al., 1993; Schulte-Merker et al., 1994). *Ntl* in the tailbud helps maintain axial progenitors by protecting them from premature differentiation induced by retinoic acid secreted by the anterior PSM and somites (Diez del Corral et al., 2003; Martin and Kimelman, 2010; Olivera-Martinez et al., 2012; Ribes et al., 2009).

Tailbud *Ntl* expression in *Greb1* morphants is indeed much lower than in wildtype or control embryos (M2: 20/34; MM: 0/11; Fig. 3Q’,R,R’). Thus, *Greb1* is required for efficient *Ntl* expression, and reduced *Ntl* levels can explain the morphant embryos’ truncated axis.

### *Greb1* depletion affects somite polarity via the segmentation clock

During axial segmentation in zebrafish embryos, a linear array of chevron-shaped somites is progressively generated from the PSM between 10-24 hpf (Fig. 3A,A’). As mentioned above, *Greb1* morphants lack morphologically discrete somites (Fig. 3A-C).

To assess if this morphological phenotype is accompanied by altered gene expression at somite boundaries, we examined *xirp2a/cb1045,* which is expressed in the myoseptum between myotomes (Deniziak et al., 2007; Schroter and Oates, 2010). Strong distinct posterior stripes of *Xirp2a* mRNA expression are frequently lost in *Greb1* morphant embryos (M1: 20/25; M2: 14/18; MM: 0/15; Fig. 3A-C), corresponding to the regions with abnormal somite appearance.

In wildtype embryos, boundaries arise between posterior and anterior compartments of adjacent somites, raising the possibility that *Greb1* is needed for somite compartmentalisation. To test this idea, we studied *myoD* transcripts, which are normally expressed in the posterior half of each somite (Weinberg et al., 1996). By contrast, expression of *myoD* extends into the anterior compartment in *Greb1* morphants (M1: 7/15; M2: 13/18; MM: 0/18; Fig. 3D-F), suggesting that anterior morphant cells have adopted a posterior character. M1 and M2 morphants show similar effects on axial morphology and *Xirp2a* and *myoD* expression, we only analysed M2 morphants in subsequent experiments.

Analysing *papC,* which is expressed in the anterior compartments of newly formed somites (Rhee et al., 2003) provides additional support for the idea that *Greb1* contributes to the establishment of anterior compartmentalisation. In morphant embryos, *papC* levels are reduced and lack clear borders (M2:14/22; MM:0/22 Fig. 3G,L).

Expression of *myoD* is normally suppressed in anterior somite compartments by *mespb* which together with *mespa,* is expressed there in newly-formed somites (Sawada et al., 2000). We examined expression of both *mesp* genes in the morphant embryos and found that, although *mespa* expression is not altered (M2: 0/24; MM:0/21 Fig. 3H,M), *mespb* expression is greatly lowered (M2: 6/10; MM: 0/21; Fig. 3 I,N). This reduction explains why *myoD* is derepressed in *Greb1* morphants, and reinforces our view that *Greb1* is needed for somite compartmentalisation.

What might cause mis-specification of somite compartments? During vertebrate axial extension, the regular production of equal-sized segments results from the action of a molecular oscillator (“segmentation clock”), which drives cyclic transcription of many PSM genes with a period corresponding to that of somite formation (Dequeant et al., 2006; Niwa et al., 2007; Palmeirim et al., 1997; Pourquie, 2011). Together, axial extension and cyclic gene expression establish reiterated expression of genes that define somite compartmentalisation and, hence, somite boundaries.

We examined two such cycling genes, *her1* and *her7*, which encode transcriptional repressors whose periodic expression in the zebrafish PSM form and pattern the somites (Oates and Ho, 2002; Pourquie, 2011; Takke and Campos-Ortega, 1999). In particular, *her7* is a regulator of *mespb* expression in forming somites (Choorapoikayil et al., 2012; Oates and Ho, 2002). Expression of *her1* is normal in the PSMs of *Greb1* morphant embryos (0/16; Fig. 3 J,O), but that of *her7* is lost, in both the tailbud and PSM (5/5, Fig. 3 K,P). The latter’s loss explains the reduced *mespb* expression and abnormal somite compartmentalisation in *Greb1* morphant embryos.

Together, our experiments support the following model for the *Greb1* morphant phenotypes (Fig. 3S). Axial extension is truncated due to reduced expression of *Ntl* and, thereby, loss of axial progenitors (Fig. 3Q,Q’,R; Martin and Kimelman, 2010), and the segmentation phenotype is caused by loss of *her7.* This model is consistent with the misregulation of *mespb* and loss of more posterior somite boundaries in both *her7* mutants and *Greb1* morphants (Figs. 3A-F; A’-F’; Oates and Ho, 2002).

As Greb1, Ntl and Her7 are all transcription factors, many of the effects on gene transcription that we observe may be direct. Greb1 doesn’t not act on the segmentation clock itself because *her1* expression still cycles (Fig. 3J,O), and low level *xirp2* expression remains (Fig. 3B,C). However, Greb1 is required for clock output via *her7,* and may also act directly on *mespb.* The latter idea would explain why *mespb* expression is abolished in the *Greb1* morphants (FIg. 3I,N). Although further experiments will be required to dissect how *Greb1* regulates gene expression in detail, the evolutionarily conserved pattern and time-course of *Greb1* expression that we have shown in mouse, chick and zebrafish (Fig. 2) suggest that *Greb1* is an important component in vertebrate axial patterning.

## MATERIAL AND METHODS

Briefly, CNH, PSM, and TBM were dissected as previously described (Fig. 1A; Cambray and Wilson, 2002), and transcription profiling carried out at the Genome Centre (Barts and the London Medical School, Blizzard Institute) using Illumina “Ref6v2” beads arrays. Microarray data were analysed within Bioconductor using the ‘lumi’ and ‘limma’ packages (Du et al., 2008; Ritchie et al., 2015). Function enrichment analysis was conducted using MetaCore (Clarivate Analytics).

We visualised spatiotemporal transcript expression in mouse, chick and zebrafish embryos by in-situ hybridisation using digoxigenin-labelled antisense RNA probes (Hanisch et al., 2013; Rallis et al., 2010; Stauber et al., 2009). To knockdown *Greb1* in zebrafish embryos, we analysed embryos which had been injected at the 1-2 cell stage with a splice-blocking or mismatch control morpholino (Fig. S1; Gene Tools, Philomath, Oregon, USA). Efficacy and specificity were tested by sizing and sequencing RT-PCR products of total RNA from morpholino-injected embryos. Further details are presented in the online Supplementary Information.

## ACKNOWLEDGEMENTS

We should like to thank the CR-UK Developmental Genetics Laboratory member, and Andrea Streit for comments on the paper. Val Wilson provided advice and training, generous help with CNH dissections, and key criticisms of a previous draft of the paper. The work was funded by Cancer Research UK and University College London.

## AUTHOR CONTRIBUTIONS

R.S.P. and D.I.H. designed the project and wrote the paper. R.S.P performed most of the experiments. R.M. provided bioinformatics analysis; A.V. contributed to studying gene expression in chick and fish.

