## Supplemental Info + Fig. S1 + Table S5 for "*Greb1* is required for axial elongation and segmentation in vertebrate embryos"

### SUPPLEMENTARY MATERIAL AND METHODS

#### Maintenance and collection of embryos

E10.5 mouse embryos were collected from *CD1* and *C57Bl/6J* pregnant females (Charles River Laboratories International, Inc. UK). Fertilized chicken eggs from Henry Stewart & Co (Louth, UK) were incubated at 37°C, and embryos were staged according to Hamburger and Hamilton (Hamburger and Hamilton, 1992). Adult wild-type zebrafish were maintained at 27°C on a regular 14 h light/10 h dark cycle, and embryos were collected and staged as described by Kimmel et al. (Kimmel et al., 1995). *p53* heterozygous and homozygous mutant zebrafish embryos were obtained by crossing *p53* homozygous female to *p53* heterozygous males (Robu et al., 2007). Animals used in this study were handled by professionals meeting all the requirements of the Animals (Scientific Procedures) Act 1986.

#### Transcription profiling

To obtain CNH, TMB, PSM tissues, mouse embryos were collected at E10.5 in M2 media (Sigma Aldrich M7167), and dissections performed as previously described (Cambray and Wilson, 2002). Approximately 50 pieces of each region were pooled separately, and total RNA extracted using the RNeasy Mini Kit (Qiagen Cat. No. 74104). Before processing the RNA samples for microarray analysis, their quality was tested using the Bioanalyser - RNA 6000 Pico kit (Agilent, Cat. No. 5067-1513). Samples with RNA integrity number (RIN) 8-10 were processed for transcriptional profiling at the Genome Centre (Barts and the London Medical School, Blizzard Institute) using Illumina “Ref6v2” beads arrays. Two biological and one technical replicate were carried out for each region – CNH, TBM, and PSM.

#### Microarray data analysis

Analysis was performed using software packages developed for Bioconductor version 2.4.0 and R version 2.9.0. The Illumina dataset were processed using the probe intensity transformation (VST) and normalization (RSN) methods from the "lumi" package (Ihaka and Gentleman, 1996; Team, 2009). Differential gene expression was assessed between tissue-type replicate groups using an empirical Bayes’ t-test as implemented in the ‘limma’ package, and taking account of replicate group and batch effects (K., 2005). Three comparisons were performed: CNH vs PSM, CNH vs TBM, CNH vs Combined PSM and TBM. The resulting p-values were adjusted to control the False Discovery Rate (FDR) using the Benjamini and Hochberg method. Two lists of differentially expressed genes were produced using different thresholds: 1) All genes that exhibited FDR<0.05 in all three comparisons, or a fold change >1.5 in the same direction in all three contrasts were classified as differentially expressed. 2) "Top50": Genes were selected on the basis of FDR<0.05 and an absolute fold change ≥1.5 from the CNH vs PSM comparison, ordered by fold change, and the top 50 most-changed genes were selected and clustered using hierarchical clustering algorithm. Genes from the two lists were combined and used to perform a pathway enrichment and network analysis with MetaCore software from Clarivate Analytics.

**In situ hybridisation**

We visualised spatiotemporal expression of various genes by in-situ hybridisations using digoxigenin-labelled antisense RNA probes (Stauber et al., 2009) (Rallis et al., 2010) (Hanisch et al., 2013). In general, templates for making antisense RNA probes for *in situ* detection of *Greb1* transcripts were generated by RT-PCR of embryonic mRNA, cloning into *PCR2.1-TOPO-TA* vector (Invitrogen), linearisation using *Spe1* or *Not1,* and transcription by T3 or T7 RNA polymerase. cDNA templates for generating other antisense-RNA probes were obtained the Julian Lewis lab.

**Morpholino injection**

To knockdown *Greb1* expression in zebrafish embryos, we injected 2 nl of the following splicing-blocking morpholinos at 2-8 ng/μl in 0.4 mM $MgSO_4$, 0.6 mM $CaCl_2$, 0.7 mM KCl, 58 mM NaCl, 5 mM HEPES pH 7.6; 0.05% phenol red: (M1) 5'-GGAAGACTGTAAAAGCTCACCCTCA-3', (M2) 5'AATACTGAAATCACACCTCTCC-TCC-3' (Gene Tools; Philomath, Oregon). Control injections used a mutated M2 oligo (MM) with 5 nucleotide mismatches: 5'-AATAGTCAAATCAGACCTGTGCTCC-3'. To test for non-specific toxicity, 4 ng/μl of blocking or control morpholino was co-injected with 6 ng/μl *p53* antisense morpholino (Robu et al., 2007). Efficacy and specificity were tested by sizing and sequencing RT-PCR products of total RNA from morpholino-injected embryos, SuperScript III One-Step RT-PCR mix (Invitrogen #12574035).

### SUPPLEMENTARY FIGURE LEGENDS

**Fig. S1:** M1 and M2 morpholinos specifically knockdown *Greb1*: (A) is schematics of *greb1* exon1-3, exon 1 and 3; and 1% agarose gel that shows wildtype and mis-spliced (red star) RT-PCR products. M1 morpholino targets exon2-intron2 boundary, and MM is control morpholino (5-nucleotide mis-matched of M2) chromatogram was obtained from Sanger sequencing for mis-spliced product-deleted exon2. B same as A and B but for M2 morpholino that targets exon16-intron16 boundary.

#### Supplementary Table Legends

Table S1: A list of differentially expressed genes. Tab1 pairwise comparisons of CNH vs PSM, and CNH vs TMB. Tab2 differentially expressed gene in CNH vs TBM comparison.

Table S2: The mouse genome informatics (MGI) data show expression of differentially expressed genes in various tissues. Tab1 list of differentially expressed genes and the annotated tissues where these genes express. Green colour show CNH up regulated and red colour CNH down regulated genes. Tab2 and tab3 graphical representation differentially expression genes in various tissues, it is an output from MGI.

Table S3: A comparison of CNH enriched genes with published CNH/NMP transcriptome. Tab1 comparison among CNH and published CNH and NMP data. Tab2 notochord genes which are also enriched in CNH.

Table S4: Results of pathway enrichment analysis (Supplementary Methods) showing the top ten enriched pathways and their associated genes.

**Table S5. List of primers sequences**

| Primer name | Sequences (5' to 3') |
|---|---|
| mGreb1 For | GCCACGGGGCGTCCGGCCCTTTC |
| mGreb1 Rev | ACCGCGCTGTGCAGGCGGGGGA |
| chGreb1-For | ATCCGCAAGGGGAGTCTTTACC-3 |
| chGreb1-Rev | GGTGAGGAGGATGAGGAGGTGA |
| zGreb1-For | AAGGAGCCACCCCTCTGCACATTCT |
| zGreb1-Rev | TTAGACGAAACCGCATTCGTCCTC |
| M1-RT-PCR-for | GGAGTCTGACCGCCAGTGACCAG |
| M1-RT-PCR-rev | AAGTGCATTACGTCCACATTCATCG |
| M2-RT-PCR-for | GCTTGTCTCTGAAGGAGGCTGAGCA |
| M2-RT-PCR-rev | ATTCTCCCTGTGGATCCATGCCAGT |
